## Supplementary Information for "Functional Connectivity-based Attractor Dynamics in Rest, Task, and Disease"

### Supplementary Figures

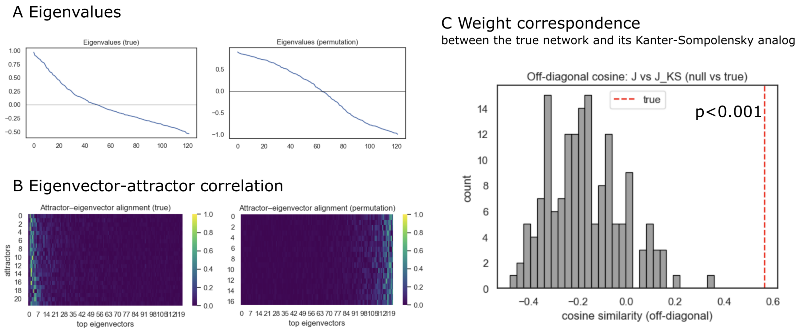


**Figure 1: Eigenstructure and projector tests of the fcANN**
**A**: Eigenvalue spectra of the empirical coupling matrix J (left) and null model 1 (coupling matrix based on phase randomized timeseries data, recalculated for each permutation) (right). **B**: Eigenvector–attractor alignment calculated from the empirical (left) and phase-randomized data (right). Attractors were obtained by deterministic relaxation from random initial states (with collapsing sign-duplicates); alignment is the absolute cosine between collapsed attractor vectors and the top eigenvectors of J. **C**: Weight correspondence between J and its Kanter–Sompolinsky (K–S) analog. From the measured attractors we formed Σ (columns are attractors) and C=$\Sigma^{T}\Sigma$/N, then computed the pseudo-inverse projector $J_{KS} = (1/N)\Sigma C^{-1}\Sigma^{T}$. Similarity was quantified as the cosine between the off-diagonal elements of J and $J_{KS}$. The gray histogram shows the null distribution from null model 2 (symmetry-preserving permutations of J, but see Source notebook for similar results with null model 1, i.e. phase randomized timeseries data); for each of the 1000 permutations $p$ we recomputed the own attractors of the surrogate network and $J_{KS}^{(p)}$. The red dashed line marks the empirical value; the one-sided p-value is the fraction of null cosines ≥ the empirical cosine. The empirical network shows stronger eigenvector–attractor alignment and substantially higher J↔J_KS off-diagonal correspondence than the null, consistent with approximate K–S projector behavior.

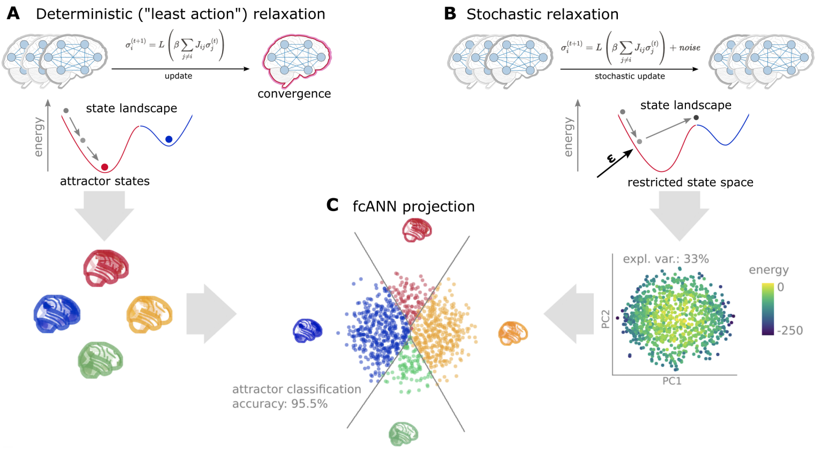


**Figure 2: Schematic representation of the fcANN projection.** The fcANN projection is a 2-dimensional visualization of the fcANN dynamics, based on the first two principal components (PCs) of the states sampled from the stochastic relaxation procedure. The first two PCs yield a clear separation of the attractor states, with the two symmetric pairs of attractor states located at the extremes of the first and second PC. To map the attractors' basins on the space spanned by the first two PCs, we obtained the attractor state of each point visited during the stochastic relaxation and fit a multinomial logistic regression model to predict the attractor state from the first two PCs. The resulting model accurately predicted attractor states of arbitrary brain activity patterns, achieving a cross-validated accuracy of 96.5% (permutation-based p<0.001). The attractor basins were visualized by using the decision boundaries obtained from this model. We propose the 2-dimensional fcANN projection as a simplified visual representation of brain dynamics, and use it as a basis for all subsequent analyses in this work.

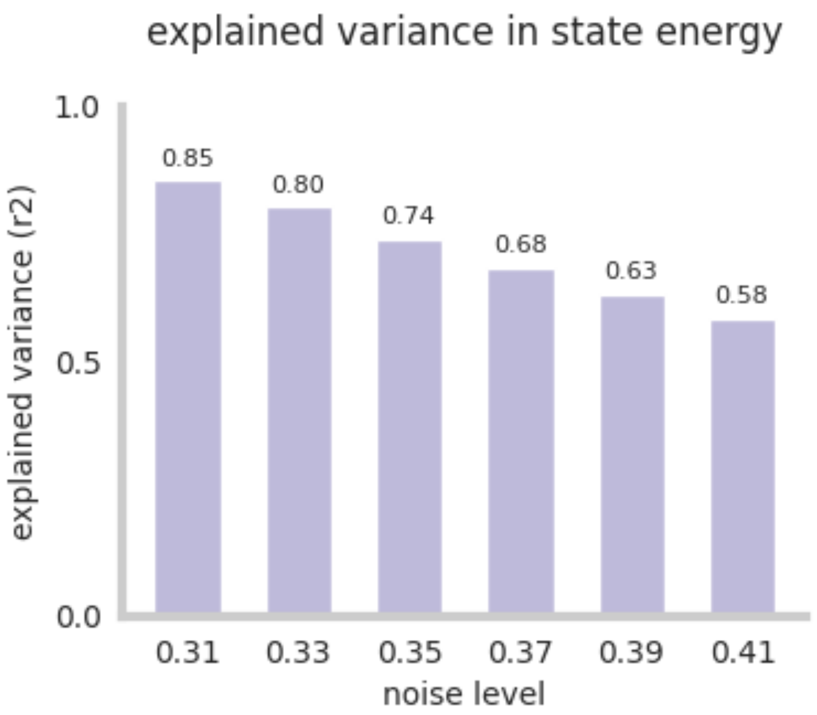


**Figure 3: Explained variance in state energy by first two principal components.** See [supplemental _material.ipynb] for details.

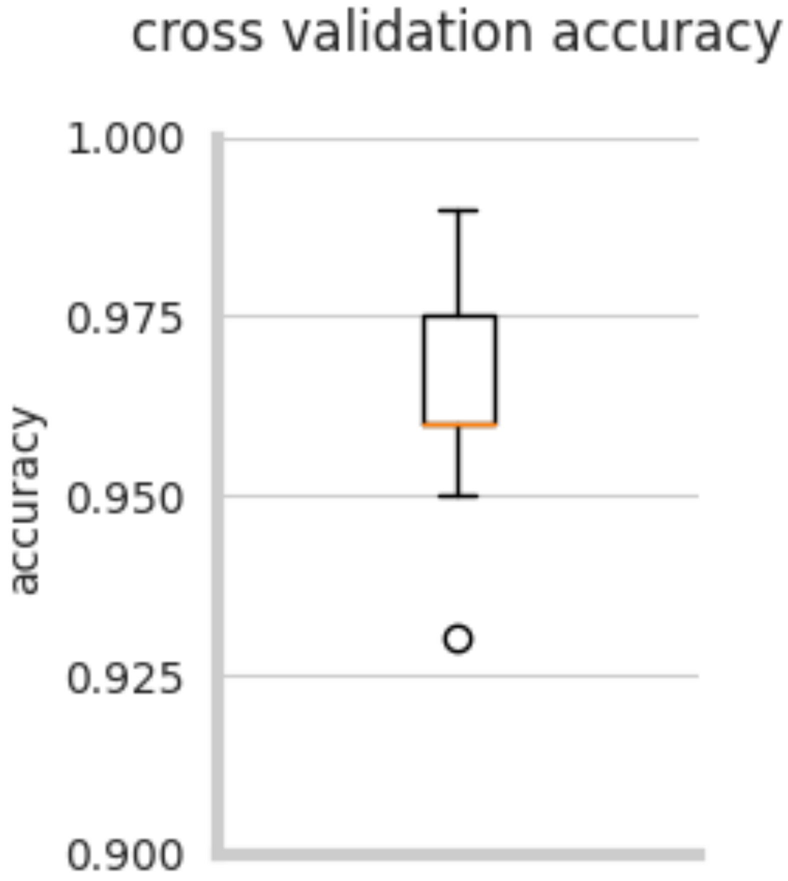

**Figure 4: Cross-validation classification accuracy of the fcANN, when predicting the attractor state from state activation.** See supplemental_material.ipynb for details.

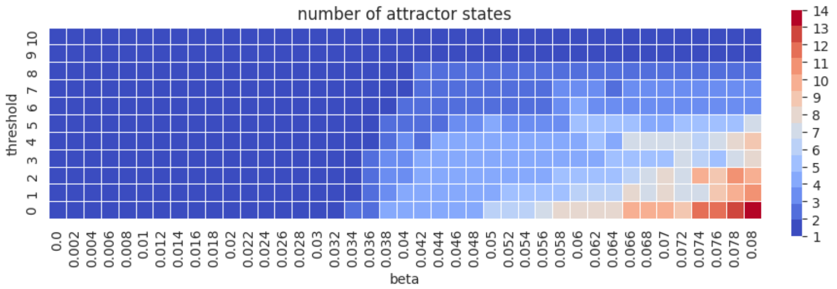


**Figure 5: Parameter sweep of fcANN parameters threshold and beta.** The number of attractor states is color-coded. See supplemental_material.ipynb for details.

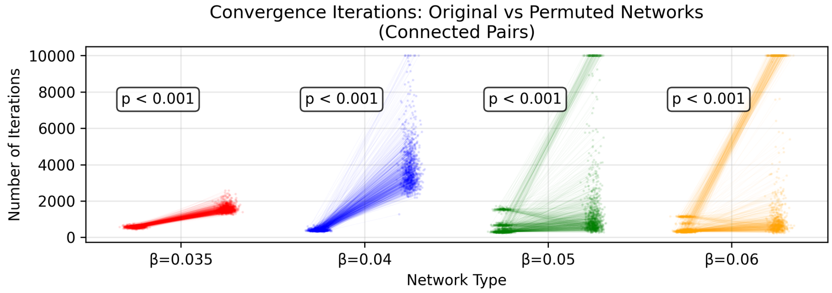


**Figure 6: fcANNs initialized with the empirical connectome have better convergence properties than permutation-based null models.** We investigated the convergence properties of functional connectome-based ANNs in study 1 by contrasting the number of iterations until reaching convergence to a permutation-based null model. In more detail, the null model was constructed by randomly permuting the upper triangle of the original connectome and filling up the lower triangle to get a symmetric network (symmetry of the weight matrix is a general requirement for convergence). This procedure was repeated 1000 times. In each repetition, we initialized both the original and the permuted fcANN with the same random input and counted the number of iterations until convergence. Each point on the plot shows an iteration number; the lines connect iteration numbers corresponding to the original and permuted matrices initialized with the same input. Statistical significance of the faster convergence in the empirical connectome was assessed via a one-sided Wilcoxon signed-rank test (i.e., a non-parametric paired test) on the paired iteration values (1,000 pairs), with the null hypothesis that the empirical connectome converges in fewer iterations than the permuted connectome. The whole procedure was repeated with $\beta = 0.3, 0.35, 0.4, 0.5$ and $0.6$ (providing 2–8 attractor states). See convergence-analysis.ipynb for details.

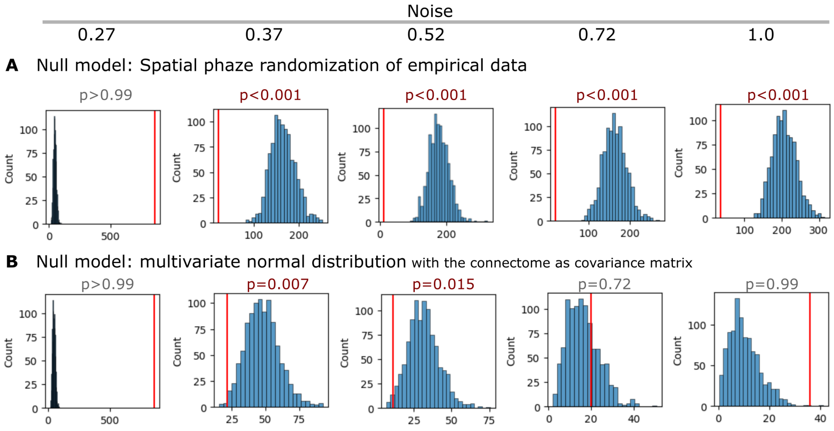


**Figure 7: Statistical inference of the fcANN state occupancy prediction with different null models. A** Results with a spatial autocorrelation-preserving null model for the empirical activity patterns. See null_models.ipynb for more details. **B** Results where simulated samples are randomly sampled from a

multivariate normal distribution, with the functional connectome as the covariance matrix, and compared to the fcANN performance. See supplemental_material.ipynb for details.

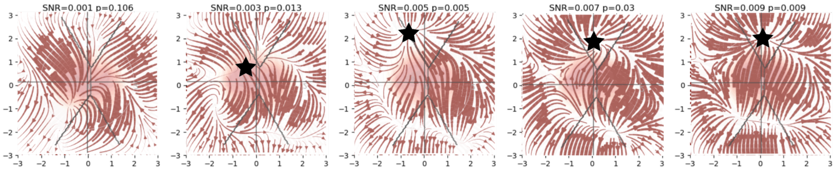


**Figure 8: FcANN can reconstruct the pain "ghost attractor".** Signal-to-noise values range from 0.003 to 0.009. Asterisk denotes the location of the simulated "ghost attractor". P-values are based on permutation testing, by randomly changing the conditions in a per-participant basis. See main_analyses.ipynb for more details.

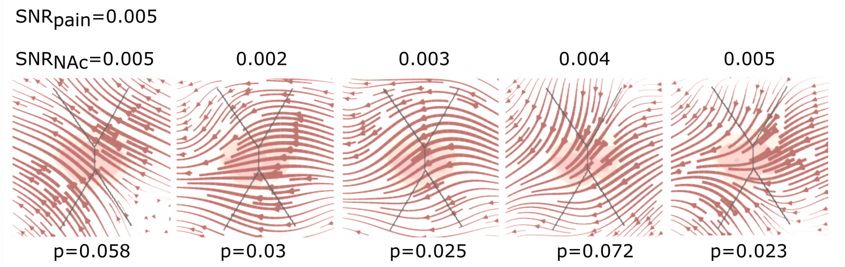


**Figure 9: fcANN can reconstruct the changes in brain dynamics caused by the voluntary downregulation of pain (as contrasted to upregulation)** Signal-to-noise values range from 0.001 to 0.005. P-values are based on permutation testing, by randomly changing the conditions in a per-participant basis. See main_analyses.ipynb for more details.

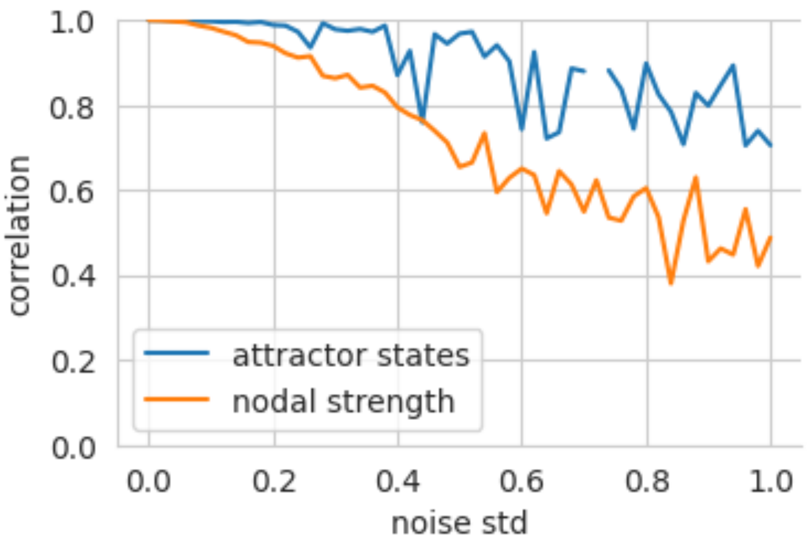


**Figure 10: Robustness of the fcANN weights to noise.** We set the temperature of the fcANN, so that two attractor states emerge and iteratively add noise to the connectome. To account for the change in dynamics, we adjust the temperature (beta) of the noisy fcANN so that exactly two states emerge. We then highlight the decrease in nodal strength of the noisy connectome (the fcANN weights) as a reference metric vs the correlation of the attractor states that emerge from the noisy connectome. See supplemental_material.ipynb for details.

| label | state | region | effect size | p value |
|---|---|---|---|---|
| 4 | execution | POSTERIOR_CINGULATE_CORTEX | 0.109 | <0.0001 |
| 3 | execution | INFERIOR_MARGINAL_SULCUS | 0.098 | <0.0001 |
| 2 | execution | CINGULATE_SULCUS_posterior | 0.092 | <0.0001 |
| 1 | execution | SOMATOMOTOR_NETWORK_mediolateral | 0.08 | 0.00976 |
| 0 | execution | SOMATOMOTOR_NETWORK_medial | 0.074 | 0.01952 |
| 11 | internal | HESCHLS GYRUS | -0.126 | <0.0001 |
| 9 | internal | SOMATOMOTOR_NETWORK_mediolateral | -0.099 | 0.00976 |
| 12 | internal | SUPERIOR_TEMPORAL_GYRUS_middle | -0.098 | <0.0001 |
| 10 | internal | FRONTAL_EYE_FIELD | -0.095 | <0.0001 |
| 7 | internal | left_SOMATOMOTOR_NETWORK_dorsolateral | -0.094 | 0.00976 |
| 5 | internal | PERIGENUAL_ANTERIOR_CINGULATE_CORTEX | -0.087 | 0.03904 |
| 8 | internal | SOMATOMOTOR_NETWORK_medial | -0.082 | 0.01952 |
| 13 | internal | LATERAL_VISUAL_NETWORK_ventroposterior | 0.077 | 0.02928 |
| 6 | internal | SUPERIOR_TEMPORAL_GYRUS_anterior | -0.069 | 0.02928 |
| 21 | external | left_MIDDLE_FRONTAL_GYRUS_posterocaudal | -0.092 | <0.0001 |
| 18 | external | POSTERIOR_CINGULATE_CORTEX | -0.089 | <0.0001 |
| 15 | external | CEREBELLUM_VIIb_medial | -0.084 | <0.0001 |
| 28 | external | INFERIOR_TEMPORAL_GYRUS | -0.073 | <0.0001 |
| 20 | external | PRECUNEUS_ventral | -0.067 | <0.0001 |
| 24 | external | left_INFERIOR_FRONTAL_SULCUS | -0.062 | <0.0001 |
| 26 | external | DORSOMEDIAL_PREFRONTAL_CORTEX_posterior | -0.06 | <0.0001 |
| 29 | external | SUPERIOR_TEMPORAL_GYRUS_middle | 0.058 | <0.0001 |
| 25 | external | right_SUPERIOR_FRONTAL_SULCUS | -0.058 | <0.0001 |
| 22 | external | left_MIDDLE_FRONTAL_GYRUS_posterorostral | -0.058 | <0.0001 |
| 30 | external | MEDIAL_VISUAL_NETWORK_anterodorsal | 0.056 | <0.0001 |
| 16 | external | PARIETO_OCCIPITAL_SULCUS_ventral | 0.052 | <0.0001 |
| 17 | external | left_MIDDLE_TEMPORAL_GYRUS_posterior | -0.051 | <0.0001 |
| 23 | external | right_MIDDLE_FRONTAL_GYRUS_posterior | -0.051 | <0.0001 |
| 14 | external | left_CEREBELLUM_CRUSII_anterior | -0.048 | 0.00976 |
| 19 | external | left_INFERIOR_PARIETAL_LOBULE | -0.047 | 0.03904 |
| 27 | external | CAUDATE_NUCLEUS_HEAD_and_NUCLEUS_ACCUMBENS | -0.043 | 0.00976 |
| 32 | perception | CEREBELLUM_VIIb_medial | 0.104 | <0.0001 |
| 43 | perception | INFERIOR_TEMPORAL_GYRUS | 0.091 | <0.0001 |
| 37 | perception | left_MIDDLE_FRONTAL_GYRUS_posterocaudal | 0.088 | <0.0001 |
| 34 | perception | POSTERIOR_CINGULATE_CORTEX | 0.08 | <0.0001 |
| 40 | perception | left_INFERIOR_FRONTAL_SULCUS | 0.066 | <0.0001 |
| 45 | perception | MEDIAL_VISUAL_NETWORK_anterodorsal | -0.065 | <0.0001 |
| 36 | perception | PRECUNEUS_ventral | 0.064 | <0.0001 |
| 38 | perception | left_MIDDLE_FRONTAL_GYRUS_posterorostral | 0.063 | <0.0001 |
| 33 | perception | left_MIDDLE_TEMPORAL_GYRUS_posterior | 0.05 | <0.0001 |
| 44 | perception | MEDIODORSAL_VISUAL_NETWORK_posterior | -0.049 | 0.02928 |
| 31 | perception | left_CEREBELLUM_CRUSII_anterior | 0.048 | 0.02928 |
| 41 | perception | right_SUPERIOR_FRONTAL_SULCUS | 0.046 | <0.0001 |
| 39 | perception | right_MIDDLE_FRONTAL_GYRUS_posterior | 0.046 | 0.02928 |
| 35 | perception | left_INFERIOR_PARIETAL_LOBULE | 0.045 | <0.0001 |
| 42 | perception | CAUDATE_NUCLEUS_HEAD_and_NUCLEUS_ACCUMBENS | 0.044 | <0.0001 |

**Figure 11: All significant differences of the mean state activation analysis on the ABIDE dataset; label denotes the region in the BASC122 atlas.** See supplemental_material.ipynb for details.

### Supplementary Tables

**Table 1: MRI acquisition parameters.** TR: repetition time; TE: echo time; FA: flip angle; FOV: field of view; EPI: echo-planar imaging; SPGR: spoiled gradient recall; SENSE/GRAPPA/ASSET: parallel imaging factors. Study 5-7 are metaanalyses or multi-center studies with varying data. Sequence parameters for these studies are available in the respective publications.

| **Parameter** | **Study 1** | **Study 2** | **Study 3** | **Study 4** |
|---|---|---|---|---|
| Scanner / head coil | Philips Achieva X 3T; 32-ch | Siemens Magnetom Skyra 3T; 32-ch | GE Discovery MR750w 3T; 20-ch | Philips Achieva TX 3T; head coil per site |
| Anatomical sequence | T1 MPRAGE | T1 MPRAGE | T1 3D IR-FSPGR | T1 SPGR (high-resolution) |
| Anatomical TR / TE | 8500 ms / 3.9 ms | 2300 ms / 2.07 ms | 5.3 ms / 2.1 ms | — / — |
| Anatomical resolution / FOV | 1×1×1 mm³; 256×256×220 mm³ | 1×1×1 mm³; 256×256×192 mm³ | 1×1×1 mm³; 256×256×172 | — |
| Resting-state EPI TR / TE / FA | 2500 ms / 35 ms / 90° | 2520 ms / 35 ms / 90° | 2500 ms / 27 ms / 81° | 2000 ms / 20 ms / — |
| Phase enc. | COL | A>>P | A>>P | — |

| | | | | |
|---|---|---|---|---|
| FOV (voxels × slices) | 240×240×132; 40 slices | 230×230×132; 38 slices | 96×96×44; 44 slices | 64×64; 42 slices |
| Slice thickness / gap / order | 3 mm / 0.3 mm / interleaved | 3 mm / 0.48 mm / interleaved | 3 mm / 0 mm / interleaved | 3 mm / — / interleaved |
| Acceleration / fat suppression | SENSE 3× / SPIR | GRAPPA 2× / Fat sat. | ASSET 2× / Fat sat. | SENSE 1.5× / — |
| Volumes / dummies / scan time | 200 / 5 / 8 min 37 s | 290 / 5 / 12 min 11 s | 240 / 0 / 10 min | — / — / — |

**Table 2: Neurosynth meta-analyses.** The table includes details about the term used for the automated meta-analyses, as well as the number of studies included in the meta-analysis, the total number of reported activations and the maximal Z-statistic from the meta-analysis.

| search term | num. studies | num. activations | max. Z |
|---|---|---|---|
| pain | 516 | 23295 | 14.8 |
| motor | 2565 | 109491 | 22.5 |
| auditory | 1252 | 46557 | 25.3 |
| visual | 3110 | 115726 | 15.4 |
| face | 896 | 31842 | 26.8 |
| autobiographical | 143 | 7251 | 15.7 |
| theory of mind | 181 | 7761 | 15.1 |
| sentences | 356 | 13461 | 16.5 |

### Supplementary Methods

**Study 4 instructions for upregulation.** *"During this scan, we are going to ask you to try to imagine as hard as you can that the thermal stimulations are more painful than they are. Try to focus on how unpleasant the pain is, for instance, how strongly you would like to remove your arm from it. Pay attention to the burning, stinging and shooting sensations. You can use your mind to turn up the dial of the pain, much like turning up the volume dial on a stereo. As you feel the pain rise in intensity, imagine it rising faster and faster and going higher and higher. Picture your skin being held up against a glowing hot metal or fire. Think of how disturbing it is to be burned, and visualize your skin sizzling, melting and bubbling as a result of the intense heat."*

**Study 4 instructions for downregulation.** *"During this scan, we are going to ask you to try to imagine as hard as you can that the thermal stimulations are less painful than they are. Focus on the part of the sensation that is pleasantly warm, like a blanket on a cold day. You can use your mind to turn down the dial of your pain sensation, much like turning down the volume dial on a stereo. As you feel the stimulation rise, let it numb your arm, so any pain you feel simply fades away. Imagine your skin is very cool, from being outside, and think of how good the stimulation feels as it warms you up."*

### Mathematical Appendix 1

**Derivation of the joint steady-state probability of free energy minimizing attractor networks** See (Spisak & Friston, 2025) for details on the whole framework.

**Regularity assumptions**

- Existence of a non-equilibrium steady-state density $p^*$ with $\Phi=\log{p^*}$ differentiable almost everywhere on its support; stationary edge fluxes are finite.
- Measurable jump kernel with finite total update rate at each configuration $\sigma$; independent Poisson clocks as specified.
- Compact support of single-site updates $x\in\left[-1,1\right]$ so boundary terms vanish in the summation-by-parts/test-function identities used for solenoidal orthogonality.

- Sufficient smoothness to justify differentiating the reversible increment at x=\sigma_i; mixed partials commute where invoked.

### Setup: Continuous–Bernoulli kernel

We assume that the single-site conditional distribution is the Continuous–Bernoulli on $[-1,1]$ with canonical parameter

$$\kappa_i(\sigma_{-i}) = b_i + \sum_{j\neq i} J_{ij}\,\sigma_j \tag{1}$$

and density (for $x \in [-1,1]$)

$$K(x \mid \kappa) = h(x) e^{\kappa x - A(\kappa)}, \qquad x \in [-1,1] \tag{2}$$

with $h(x) = \backslash tfrac121_{[-1,1]}(x)$ and $A(\kappa) = \log(\sinh\kappa) - \log$ (so the $\kappa \to 0$ limit is uniform on $[-1,1]$).

The conditional mean is:

$$L(\kappa) = E_{K(\cdot|\kappa)}[x] = \coth\kappa - \frac{1}{\kappa} \tag{3}$$

### 1. Master equation

Let's consider a precise formalization of the asynchronous update procedure: each site $i$ has an independent Poisson clock of rate $\gamma_i$. When it rings, $\sigma_i$ is replaced by a draw $x \sim K\big(\cdot\,|\,\kappa_i(\sigma_{-i})\big)$.

With $\sigma^{(i,x)}$ the configuration equal to $\sigma$ but with coordinate $i$ set to $x$, the transition density is

$$w_i\big(\sigma^{(i,x)} \mid \sigma\big) = \gamma_i K\big(x \mid \kappa_i(\sigma_{-i})\big) \tag{4}$$

The master equation for $p(\sigma, t)$ is

$$\partial_t p(\sigma,t) = \sum_{i=1}^{N} \int_{-1}^{1} \left[ \underbrace{w_i\big(\sigma \mid \sigma^{(i,x)}\big) p\big(\sigma^{(i,x)}, t\big)}_{inflow} - \underbrace{w_i\big(\sigma^{(i,x)} \mid \sigma\big) p(\sigma,t)}_{outflow} \right] dx \tag{5}$$

> **Note**
>
> In a small interval $dt$, probability in state $\sigma$ changes by inflow from all one-site predecessors $\sigma^{(i,x)}$ minus outflow from $\sigma$ to those states. Multiple updates within $dt$ are $O(dt^2)$ and negligible. Because the updated value $x$ is continuous, we integrate over $x \in [-1,1]$.

### 2. Probability currents

Let's denote the site-wise flux density as:

$$F_i(\sigma \to x) := w_i\big(\sigma^{(i,x)} \mid \sigma\big) p^*(\sigma) \tag{6}$$

At steady state $p^*$, by definition we have:

$$\sum_i \int_{-1}^{1} \Big(F_i\big(\sigma^{(i,x)} \to \sigma\big) - F_i(\sigma \to x)\Big)\, dx = 0 \tag{7}$$

### 3. Detailed balance condition

A special (and maybe the most intuitive) way to satisfy the previous eq. is the case of detailed balance or equilibrium. In this case, every transition is balanced:

$$F_i\big(\sigma^{(i,x)} \to \sigma\big) = F_i(\sigma \to x) \tag{8}$$

It can be shown that:

$$w_i\big(\, \sigma^{(i,x)} \mid \sigma \,\big)p^*(\sigma) = w_i\big(\, \sigma \mid \sigma^{(i,x)} \,\big)p^*\big(\sigma^{(i,x)}\big) \Leftrightarrow \partial_{\sigma_i}\Phi(\sigma) = \kappa_i(\sigma_{-i}) = b_i + \sum_{k\neq i} J_{ik}\sigma_k \tag{9}$$

I.e., the log-density $\Phi$ changes by the local slope at $i$ (see detailed derivation below).

Hence for $j \neq i$:

$$\partial_{\sigma_j}\partial_{\sigma_i}\Phi = J_{ij}, \qquad \partial_{\sigma_i}\partial_{\sigma_j}\Phi = J_{ji} \tag{10}$$

Mixed partials commute, thus:

$$\textit{detailed balance} \;\Rightarrow J_{ij} = J_{ji} \tag{11}$$

Therefore, **equilibrium (detailed balance) is possible only when the coupling is symmetric**.

**Note**

**Derivation**

From:

$$w_i\big(\, \sigma^{(i,x)} \mid \sigma \,\big)p^*(\sigma) = w_i\big(\, \sigma \mid \sigma^{(i,x)} \,\big)p^*\big(\sigma^{(i,x)}\big)$$

We take logs ($\Phi = \log p^*$) and rearrange:

$$\Phi\big(\sigma^{(i,x)}\big) - \Phi(\sigma) = \log w_i\,\big(\, \sigma^{(i,x)} \mid \sigma \,\big) - \log w_i\,\big(\, \sigma \mid \sigma^{(i,x)} \,\big)$$

Substitute $w_i = \gamma_i K(\,\cdot \mid \kappa\,)$ with $\kappa = \kappa_i(\sigma_{-i}) = \kappa_i\big(\sigma_{-i}^{(i,x)}\big)$:

$$\Phi\big(\sigma^{(i,x)}\big) - \Phi(\sigma) = \backslash big[\log\gamma_i + \log K\,(\,x \mid \kappa\,)\backslash big] - \backslash big[\log\gamma_i + \log K\,(\sigma_i|\kappa)\backslash big]$$
$$= \log K\,(\,x \mid \kappa\,) - logK(\,\sigma_i \mid \kappa\,)$$

Use CB form $\log K\,(z|\kappa) = \log h\,(z) + \kappa z - A(\kappa)$:

$$\Phi\big(\sigma^{(i,x)}\big) - \Phi(\sigma) = \backslash big[\log h\,(x) - \log h\,(\sigma_i)\backslash big] + \kappa(x - \sigma_i)$$

($A(\kappa)$ cancels because $\kappa$ is the same).

For Continuous–Bernoulli on $[-1,1]$, $h(x) = 1/2$ on the support $\Rightarrow \log h\,(x) - logh(\sigma_i) = 0$, so:

$$\Phi\big(\sigma^{(i,x)}\big) - \Phi(\sigma) = \kappa(x - \sigma_i) \tag{12}$$

Differentiating at $x = \sigma_i$ yields

$$\partial_{\sigma_i}\Phi(\sigma) = \kappa_i(\sigma_{-i}) = b_i + \sum_{k\neq i} J_{ik}\sigma_k \tag{13}$$

### 4. Non-equilibrium Steady State (NESS)

Detailed balance (equilibrium) is only one specific way the steady-state condition can hold:

$$\sum_i \int_{-1}^{1} \left(F_i\left(\sigma^{(i,x)} \to \sigma\right) - F_i(\sigma \to x)\right) dx = 0 \tag{14}$$

Let's consider the general case: an arbitrary (possibly asymmetric) $J$ coupling. $J$ can always be decomposed into:

$$J = J^{\mathrm{S}} + J^{\mathrm{A}}, \qquad J^{\mathrm{S}} := \tfrac{1}{2}(J + J^{\top}), \quad J^{\mathrm{A}} := \tfrac{1}{2}(J - J^{\top}) \tag{15}$$

This leads to a decomposition of the edge currents on each directed edge $\left(\sigma, \sigma^{(i,x)}\right)$:

$$F_i^{\mathrm{sym}}(\sigma \to x) := \tfrac{1}{2}\left(F_i(\sigma \to x) + F_i\left(\sigma^{(i,x)} \to \sigma\right)\right), \qquad F_i^{\mathrm{sol}}(\sigma \to x) := \tfrac{1}{2}\left(F_i(\sigma \to x) - F_i\left(\sigma^{(i,x)} \to \sigma\right)\right) \tag{16}$$

so that $F_i = F_i^{\mathrm{sym}} + F_i^{\mathrm{sol}}$ and $F_i^{\mathrm{sym}}(\sigma \to x) = F_i^{\mathrm{sym}}\left(\sigma^{(i,x)} \to \sigma\right)$.

Since $F_i(\sigma \to x) = e^{\Phi(\sigma)} w_i(\sigma \to x)$, define rates accordingly:

$$w_i^{\mathrm{sym}}(\sigma \to x) := e^{-\Phi(\sigma)} F_i^{\mathrm{sym}}(\sigma \to x), \qquad w_i^{\mathrm{sol}}(\sigma \to x) := e^{-\Phi(\sigma)} F_i^{\mathrm{sol}}(\sigma \to x) \tag{17}$$

so $w_i = w_i^{\mathrm{sym}} + w_i^{\mathrm{sol}}$ and $F_i^{\mathrm{sym}} = e^{\Phi} w_i^{\mathrm{sym}}$, $F_i^{\mathrm{sol}} = e^{\Phi} w_i^{\mathrm{sol}}$.

We proceed in two steps: (i) the antisymmetric component is divergence-free and never changes the steady state; (ii) the remaining symmetric component yields exactly the same $p^*$ as the equilibrium distribution obtained by setting $J = J^{\mathrm{S}}$. Finally, as a bonus, we show that the induced circulating flow is indeed tangential to the level sets of $\Phi$, in accordance to the Helmholtz decomposition.

*i) Antisymmetric component is divergence-free (does not change $p^*$).*

Starting again from stationarity of the full current,

$$\sum_i \int_{-1}^{1} \left(F_i\left(\sigma^{(i,x)} \to \sigma\right) - F_i(\sigma \to x)\right) dx = 0 \tag{18}$$

subtract the same identity written with $F^{\text{sym}}$ in place of $F$. Because $F^{\text{sym}}(\sigma \to x) = F^{\text{sym}}\left(\sigma^{(i,x)} \to \sigma\right)$ on every edge, its contribution vanishes pairwise, giving

$$\sum_i \int_{-1}^{1} \left(F_i^{\text{sol}}\left(\sigma^{(i,x)} \to \sigma\right) - F_i^{\text{sol}}(\sigma \to x)\right) dx = 0 \tag{19}$$

Thus $F^{\text{sol}}$ has zero divergence and does not change $p^*$.

*ii) Symmetric component determines p* and matches the equilibrium solution with $J = J^{\text{S}}$.*

From Section 3, only $J^{\text{S}}$ can appear in a potential gradient (commuting mixed partials). Therefore

$$\partial_{\sigma_i}\Phi(\sigma) = b_i + \sum_{k \neq i} J_{ik}^{\text{S}}\ \sigma_k \tag{20}$$

which integrates to the same $\Phi = \log p^*$ you obtain by enforcing detailed balance with the symmetric coupling $J = J^{\text{S}}$ (i.e., in the absence of $J^{\text{A}}$). Equivalently, the symmetric current $F^{\text{sym}}$ is reversible with respect to $p^*$ and alone reproduces the same steady state.

*Note: Tangency to level sets of $\Phi$ (no work against the potential).*

Multiply the preceding identity by a test function $\psi(\sigma)$ and integrate over $\sigma$ (“summation by parts”) to obtain

$$\int d\sigma \sum_i \int_{-1}^{1} F_i^{\text{sol}}(\sigma \to x)\ \left[\psi\left(\sigma^{(i,x)}\right) - \psi(\sigma)\right] dx = 0 \tag{21}$$

Taking $\psi = \Phi$ yields

$$-\int d\sigma \sum_i \int_{-1}^{1} F_i^{\text{sol}}(\sigma \to x)\ \left[\Phi\left(\sigma^{(i,x)}\right) - \Phi(\sigma)\right] dx = 0 \tag{22}$$

so the solenoidal current is orthogonal to discrete gradients and flows tangent to the isocontours of $\Phi$ (the level sets of $p^*$).

1. Link to $J^{\text{A}}$.

Under the linear Continuous–Bernoulli parametrization, $J^{\text{A}}$ cannot contribute to any scalar potential (mixed partials must commute). Therefore, varying $J^{\text{A}}$ at fixed $J^{\text{S}}$ leaves $\Phi$ and $p^*$ unchanged; it only affects the circulating, divergence-free component $F^{\text{sol}}$.

*Conclusion: explicit steady-state $p^*$ for arbitrary $J$*

By the results above, only the symmetric component $J^{\mathrm{S}} := \frac{1}{2}(J + J^{\top})$ can shape the potential $\Phi = \log p^*$. Therefore, for arbitrary (possibly asymmetric) $J$, the non-equilibrium steady-state density takes the Boltzmann-like form:

$$p^*(\sigma) = \frac{1}{Z}\exp\left(\sum_i b_i\ \sigma_i + \sum_{i,j} J_{ij}^{\mathrm{S}}\ \sigma_i\,\sigma_j\right) \tag{23}$$

with $J_{ii}^{\mathrm{S}} = 0$ and partition function

$$Z = \int_{[-1,1]^N} exp\left(\sum_i b_i\ \sigma_i + \sum_{i,j} J_{ij}^{\mathrm{S}}\ \sigma_i\,\sigma_j\right) d \tag{24}$$

The antisymmetric component $J^{\mathrm{A}} := \frac{1}{2}(J - J^{\top})$ contributes only to divergence-free (solenoidal) probability currents tangent to the level sets of $\Phi$ and thus does not alter $p^*$. This expression matches the manuscript's stationary density, with $J^{\mathrm{S}}$ playing the role of the effective (symmetric) interaction matrix.

### Supplementary Information 2

Here we provide a concise derivation of how σ and $J$ changes as a consequence of free energy minimization resulting in an inference (node update) rule and a local, incremental learning rule, respectively.

For more background and a detailed derivation, see (Spisak & Friston, 2025).

Let the instantaneous net input to node $\sigma_i$ be

$$u_i := b_i + \sum_{j \neq i} J_{ij}\ \sigma_j, \tag{25}$$

and let $q(\sigma_i) \propto e^{b_q \sigma_i}$ be the variational marginal with mean $L(b_q)$, where $L$ is the Langevin function (the mean of the Continuous–Bernoulli).

#### Inference (Hopfield-style)

We start by writing up the local variational free energy from the point of view of a single node $\sigma_i$:

$$F = E_q[\ln q\,(\sigma_i)] - E_q\left[\ln p\left(\sigma_{\setminus i} \mid \sigma_i\right)\right] \tag{26}$$

Next we express:

$$\ln q\,(\sigma_i) = b_q\,\sigma_i - A(b_q) + \ln h\,(\sigma_i) \tag{27}$$

with mean $E_q[\sigma_i] = L(b_q) = A'(b_q)$.

and

$$\ln p\,(\,\sigma_{\setminus i}\mid \sigma_i\,) = \text{const} + \sigma_i\,\textstyle\sum_{j\neq i} J_{ij}\,\sigma_j. \tag{28}$$

As it depends on $\sigma_i$ only through the linear slope $\sum_{j\neq i} J_{ij}\,\sigma_j$:

$$E_q[\,\ln p\,(\,\sigma_{\setminus i}\mid \sigma_i\,)\,] = \text{const} + L(b_q)\,\textstyle\sum_{j\neq i} J_{ij}\,\sigma_j \tag{29}$$

Now, we assemble the local free energy (dropping constants independent of $b_q$):

$$F = (b_q - b_i)\,L(b_q) - [A(b_q) - A(b_i)] - L(b_q)\textstyle\sum_{j\neq i} J_{ij}\,\sigma_j. \tag{30}$$

Equivalently, using $u_i = b_i + \sum_{j\neq i} J_{ij}\,\sigma_j$ and $A'(b_q) = L(b_q)$,

$$F = (b_q - u_i)\,L(b_q) - A(b_q) + \text{const}. \tag{31}$$

1. Differentiate w.r.t. $b_q$ and use $A'(b_q) = L(b_q)$ to cancel terms:

$$\frac{\partial F}{\partial b_q} = (b_q - u_i)\,L'(b_q) + L(b_q) - A'(b_q) = (b_q - u_i)\,L'(b_q). \tag{32}$$

Setting the derivative to zero gives $b_q^{\star} = u_i$ and therefore the node-wise update

$$\boxed{E_q[\sigma_i] \;=\; L(b_q^{\star}) \;=\; L!\left(b_i + \textstyle\sum_{j\neq i} J_{ij}\;\sigma_j\right)} \tag{33}$$

which is the stochastic Hopfield/Boltzmann-style activation with the Langevin nonlinearity.

### Learning (Hebb – anti-Hebb)

Make the dependence on $u_i$ explicit by adding and subtracting the CB log-partition $A(u_i)$ (using $\ln p\left(\sigma_i \mid \sigma_{\setminus i}\right) = u_i\,\sigma_i - A(u_i) + \ln h\,(\sigma_i)$ and Bayes' rule):

$$F = E_q\left[\left(b_q - u_i\right)\sigma_i\right] + A(u_i) - A\left(b_q\right) + \text{const.} \tag{34}$$

Now

$$\frac{\partial F}{\partial u_i} = -E_q[\sigma_i] + A'(u_i) = -E_q[\sigma_i] + L(u_i), \tag{35}$$

and by the chain rule with $\partial u_i / \partial J_{ij} = \sigma_j$ we obtain

$$\frac{\partial F}{\partial J_{ij}} = \left[\,L(u_i) - E_q[\sigma_i]\,\right]\sigma_j. \tag{36}$$

Using a stochastic (sample-based) estimate $E_q[\sigma_i] \approx \sigma_i$ and descending $F$ gives the local update

$$\boxed{\Delta J_{ij} \;\propto\; \underbrace{\sigma_i\sigma_j}_{\text{Hebbian (observed)}} \;-\; \underbrace{L!\,(b_i + \textstyle\sum_{k\neq i} J_{ik}\,\sigma_k)\sigma_j}_{\text{anti-Hebbian (predicted)}}} \tag{37}$$

This is a predictive coding-based learning rule that strengthens observed correlations and subtracts those already predicted by the current model.

### Mathematical Appendix 2

**Self-orthogonalization of attractor states**

To illustrate how the learning rule in free energy minimizing attractor networks gives rise to efficient, (approximately) orthogonal representations of the external states, suppose the network has already learned a pattern $s^{(1)}$, whose neural representation is the attractor $\sigma^{(1)}$ and associated weights $J^{(1)}$. When a new pattern $s^{(2)}$ is presented that is correlated with $s^{(1)}$, the network's prediction for $\sigma_i^{(2)}$ will be $\hat{\sigma}_i = L\left(b_i + \sum_{k\neq i} J_{ik}^{(1)}\sigma_k\right)$. Because inference with $J^{(1)}$ converges to $\sigma^{(1)}$ and $\sigma^{(2)}$ is correlated with $\sigma^{(1)}$, the prediction $\hat{\sigma}$ will capture variance in $\sigma^{(2)}$ that is 'explained' by $\sigma^{(1)}$. The learning rule updates the weights based only on the unexplained (residual) component of the variance, the prediction error. In other words, $\hat{\sigma}$ approximates the projection of $\sigma^{(2)}$ onto the subspace already spanned by $\sigma^{(1)}$. Therefore, the weight update primarily strengthens weights corresponding to the component of $\sigma^{(2)}$ that is orthogonal to $\sigma^{(1)}$. Thus, the learning effectively encodes this residual, $\sigma_{\perp}^{(2)}$, ensuring that the new attractor components being formed tend to be orthogonal to those already established.

For more details on self-orthogonalization in these networks, including an empirical demonstration, see (Spisak & Friston, 2025).

### Mathematical Appendix 3

**Reconstruction of the attractor network from the activation timeseries.**

We start from Eq. (1), written in matrix notation: $E_{HN}(\sigma) = -\frac{1}{2}\sigma^\top J\sigma + \sigma^\top b\backslash= -\frac{1}{2}\sigma^\top J\sigma + \sigma^\top JJ^{-1}b\backslash= -\frac{1}{2}\sigma^\top J\sigma + \sigma^\top JJ^{-1}b - \frac{1}{2}b^\top J^{-1}b + \frac{1}{2}b^\top J^{-1}b$

To complete the square, we added and subtracted $\frac{1}{2}b^\top J^{-1}b$ within the exponent. We recognize that $\sigma^\top JJ^{-1}b = \sigma^\top b$. Now add and subtract $\frac{1}{2}b^\top J^{-1}b$

$= -\frac{1}{2}[\sigma^\top J\sigma - 2\sigma^\top JJ^{-1}b + b^\top J^{-1}b] + \frac{1}{2}b^\top J^{-1}b\backslash= -\frac{1}{2}[\sigma - J^{-1}b]^\top J[\sigma - J^{-1}b] + \frac{1}{2}b^T J^{-1}b$

Given that $P(\sigma) = \exp\left(-E_{HN}(\sigma)\right)$, the expression simplifies to: $P(\sigma) \propto \exp\left(-\frac{1}{2}(\sigma - J^{-1}b)^\top J(\sigma - J^{-1}b)\right)$

Note that the term $\frac{1}{2}b^\top J^{-1}b$ is independent of $\sigma$; we have absorbed it into the normalization constant. This is exactly the exponent of a multivariate Gaussian distribution with mean $J^{-1}b$ and covariance matrix $J^{-1}$, meaning that the weight matrix of the attractor network can be reconstructed as the inverse covariance matrix of activation timeseries of the lower-level nodes: $J = -\Lambda = -\Sigma^{-1}$
